# Lmo7 recruits myosin II heavy chain to induce apical constriction in *Xenopus* ectoderm

**DOI:** 10.1101/2021.05.12.443820

**Authors:** Miho Matsuda, Chih-Wen Chu, Sergei Y. Sokol

## Abstract

Apical constriction, or reduction of the apical domain, underlies many morphogenetic events during development, such as furrow or tube formation. Actomyosin complexes play an essential role in apical constriction, however the detailed analysis of molecular mechanisms is still pending. Here we show that Lim domain only protein 7 (Lmo7), a multidomain adaptor at apical junctions, promotes apical constriction in the *Xenopus* superficial ectoderm, whereas apical domain size increases in Lmo7-depleted cells. Lmo7 is localized to and promotes the formation of the circumferential actomyosin belt adjacent to apical junctions. We find that Lmo7-dependent apical constriction requires the RhoA-ROCK-Non-muscle myosin II (NMII) pathway. Strikingly, Lmo7 binds and recruits NMII heavy chains to apical junctions. Lmo7 overexpression altered the subcellular distribution of Wtip, a sensor of mechanical tension. Our findings suggest that Lmo7 serves as a scaffold organizing the actomyosin network to regulate contractility at apical junctions.

## Introduction

Apical constriction, or reduction of apical domain, underlies a variety of morphogenetic processes during embryonic development (Martin and Goldstein, 2014; Perez-Vale and Peifer, 2020; Sawyer et al., 2010). One consequence of apical constriction is cell delamination required for epithelial-to-mesenchymal transition (EMT)(Ramkumar et al., 2016; Williams et al., 2012). Another consequence is epithelial invagination that leads to epithelial furrow or tube formation (Eiraku et al., 2011; Sherrard et al., 2010; Wallingford et al., 2013). Significantly, failure of apical constriction results in epithelial morphogenesis defects, such as abnormal neural tube closure (Balashova et al., 2017; Brown and Garcia-Garcia, 2018; Colas and Schoenwolf, 2001; Haigo et al., 2003; Itoh et al., 2014; Kowalczyk et al., 2021).

The driving force for apical constriction is contractions of actomyosin networks associated with apical junctions, which are mainly composed of adherens junctions (AJs) and tight junctions (TJs)(Coravos and Martin, 2016; Gorfinkiel and Blanchard, 2011; Munjal et al., 2015). The actomyosin system is a highly dynamic network of filamentous actin (F-actin) and non-muscle myosin II (NMII), crosslinked by *α*- actinin (Vicente-Manzanares et al., 2009). NMII heavy chains assemble tail-to-tail into bipolar filaments that form NMII mini-filaments (Bresnick, 1999; Heissler and Sellers, 2014; Shutova and Svitkina, 2018). The head domains of NMII mini-filaments bind F-actin and the sliding movements between F-actin and NMII generate contractile forces mediating apical constriction (Lecuit et al., 2011; Martin and Goldstein, 2014; Salbreux et al., 2012). Actomyosin complexes can be arranged in many patterns from loose mesh-like networks to highly organized sarcomere-like actomyosin bundles (Agarwal and Zaidel-Bar, 2019; Schwayer et al., 2016). Epithelial cells with constricted apical domains contain highly organized perijunctional actomyosin bundles (Boller et al., 1985; Hirano et al., 1987; Yonemura et al., 1995). These actomyosin bundles are linked to apical junctions by F-actin binding proteins, including *α*E-catenin, vinculin and afadin at AJs, and zonule occludens proteins (ZO-1-3) at TJs (Abe and Takeichi, 2008; Drees et al., 2005; Fanning et al., 1998; Itoh et al., 1997; Sawyer et al., 2009; Yamada et al., 2005). Uncoupling of actomyosin from apical junctions as observed in *canoe/afadin* deficient *Drosophila* embryos leads to failure of apical constriction during morphogenesis (Sawyer et al., 2011; Sawyer et al., 2009).

One common mechanism of apical constriction is the recruitment and activation of Rho-associated kinase (ROCK) at apical junctions. The GTP-bound form of Rho GTPase (Rho-GTP) binds and activates ROCK which phosphorylates myosin regulatory light chain (MRLC) needed for NMII conformation change and bipolar filament formation (Amano et al., 1996). For instance, Rho guanine nucleotide exchange factors (RhoGEFs), such as p114RhoGEF, GEF-H1 and Plekhg5, accelerate the intrinsic exchange activity of Rho GTPase to promote the formation of Rho-GTP (Kolsch et al., 2007; Popov et al., 2018; Terry et al., 2011). Epb41l4 and Epb41l5 bind and activate p114RhoGEF at apical junctions (Nakajima and Tanoue, 2011; Nakajima and Tanoue, 2012). Shroom3 directly binds and recruits ROCK to apical junctions (Das et al., 2014; Nishimura and Takeichi, 2008). Depletion of either GEF-H1, Plekhg5, Epb41l5 or Shroom3 inhibits apical constriction in various models. For instance, neural tube (NT) failed to close in *shroom3* null or *gef-h1* knockdown embryos (Haigo et al., 2003; Hildebrand, 2005; Hildebrand and Soriano, 1999; Itoh et al., 2014). Blastopore lip formation failed in *plekhg5* knockdown embryos during gastrulation (Popov et al., 2018). In *epb41l5* null embryos E-cadherin is stabilized at AJs, leading to incomplete EMT during mesoderm formation (Lee et al., 2007). These findings confirm the importance of Rho-ROCK-NMII signaling and actomyosin contractility in apical constriction during epithelial morphogenesis.

To gain further insights into the interaction of the actomyosin network with apical junctional machinery, we have initiated the study of LIM domain only protein 7 (Lmo7) that was originally identified as a binding partner of Afadin and *α*-actinin (Ooshio et al., 2004). Lmo7 has been implicated in skeletal and cardiac muscle differentiation and in metastatic cancer (He et al., 2018; Holaska et al., 2006; Liu et al., 2021; Nakamura et al., 2011; Ott et al., 2008). The *Drosophila* homologue Smallish is essential for epithelial morphogenesis and controls apical domain size in the follicular epithelium (Beati et al., 2018). On the other hand, mice lacking *lmo7* are viable and develop relatively minor defects (Du et al., 2019; Lao et al., 2015; Mull et al., 2015; Tanaka-Okamoto et al., 2009). These observations suggest that Lmo7 may have uncharacterized functions in the reorganization of apical junctions and actomyosin in vertebrate epithelia.

Our study describes Lmo7 as a novel regulator of apical constriction in *Xenopus* ectoderm cells. This activity of Lmo7 requires the Rho-ROCK-NMII system, however the primary mechanism appears to be the binding and recruitment of NMII heavy chain to actomyosin bundles that are proximal to apical junctions. Lmo7 also stimulates F-actin accumulation at apical junctions to generate thicker and denser actomyosin bundles, although this accumulation of F-actin may not be required to induce apical constriction in individual cells. We propose that Lmo7 is a scaffold protein that regulates crosstalk between apical junctions and actomyosin contractility.

## Results

### Lmo7 is localized in the proximity of apical cell junctions in *Xenopus* ectoderm

To understand Lmo7 function in epithelial morphogenesis, we studied Lmo7 protein distribution in *Xenopus* ectoderm, a tissue that exhibits dynamic remodeling of apical junctions (Higashi et al., 2016). In ectoderm cells of stage 11 embryos, Lmo7 localized differently from other components of apical junctions. First, Lmo7 formed two distinct parallel bands along cell-cell boundaries, especially in short cell contact areas (Fig. 1A, B). By contrast, ZO-1, a TJ component (Hartsock and Nelson, 2008), was detected as a single band flanked by two Lmo7 bands (Fig. 1C, D). Also, Lmo7 accumulated near but not at tricellular junctions (Fig. 1B’). These observations reveal Lmo7 in the close proximity of apical junctions.

**Figure 1.**
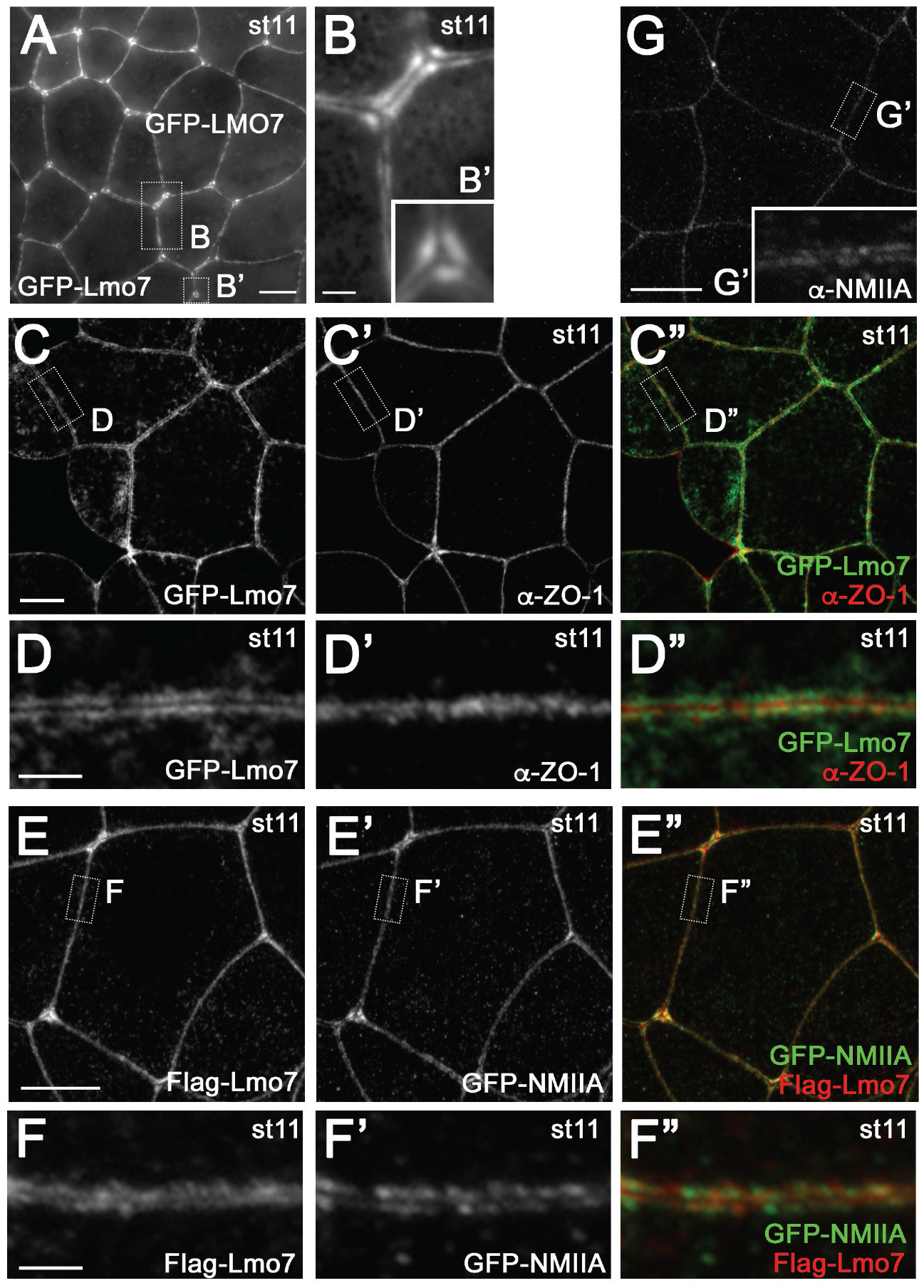
Lmo7 localizes at and near junctional actomyosin bundles in *Xenopus* ectoderm cells. *Xenopus* embryos were injected into animal blastomeres with different mRNAs and the superficial ectoderm was imaged at stage 11. Protein localization was evaluated by direct GFP fluorescence of GFP- Lmo7 and GFP-NMIIA, or indirect immunofluorescence for Flag-Lmo7, endogenous NMIIA and ZO-1. (A) Embryos injected with GFP-Lmo7 mRNA (200 pg). Areas marked by rectangles in A are enlarged in B and B’. (B) Peri-junctional double bands of GFP-Lmo7 are more evident in shorter cell-cell junctions. (B’) GFP-Lmo7 is enriched near tricellular junctions but excluded from them. (C-C”) Relative distribution of GFP-Lmo7 and endogenous ZO-1 in embryos injected with GFP-Lmo7 mRNA (500 pg). Areas marked by rectangles in C-C” are enlarged in D-D”. (D-D”) ZO-1 single band is positioned between GFP-Lmo7 double bands. (E-E”) Relative protein localization in embryos injected with Flag- Lmo7 (200 pg) and GFP-NMIIA (300 pg) mRNAs. Areas marked by rectangles in E-E” are enlarged in F-F”. (F-F”) Flag-Lmo7 and GFP-NMIIA double bands largely overlap, although Flag-Lmo7 locates slightly closer to the cell-cell border. (G-G’) Endogenous NMIIA localization. An area marked by a rectangle is enlarged in G’. (G’) NMIIA forms double bands along the cell-cell boundary. Scale bars: 10 μm in A, C, E and G. 2 μm in B, B’, D and F. Images shown are representative of two to ten experiments.

We next asked whether the location of Lmo7 corresponds to the apical circumferential actomyosin bundles that undercoat apical junctions (Yonemura et al., 1995). The major components of actomyosin bundles are F-actin, NMII and *α*-actinin, an F-actin crosslinker (Murphy and Young, 2015). We observed that both endogenous NMIIA and exogenous GFP-NMIIA form two bands along cell-cell boundaries in the *Xenopus* superficial ectoderm (Fig. 1E’, F’, G-G’). The GFP-NMIIA bands largely overlapped with those of Flag-Lmo7 (Fig. 1E-F”). These results suggest that Lmo7 localizes at or near apical junctional actomyosin bundles.

### Lmo7 induces apical constriction in *Xenopus* superficial ectoderm

A previous study showed that Smallish, the Lmo7 homologue in *Drosophila*, reduces apical domain in follicular epithelial cells (Beati et al., 2018). We therefore asked whether Lmo7 has a similar activity that could be evaluated by the enrichment of apical pigment granules in *Xenopus* ectoderm (Haigo et al., 2003; Itoh et al., 2014; Merriam et al., 1983; Morita et al., 2010; Ossipova et al., 2014). Injection of GFP-Lmo7 RNA triggered pigment granule accumulation accompanied by epithelial invagination (Fig. 2A, B).

**Figure 2.**
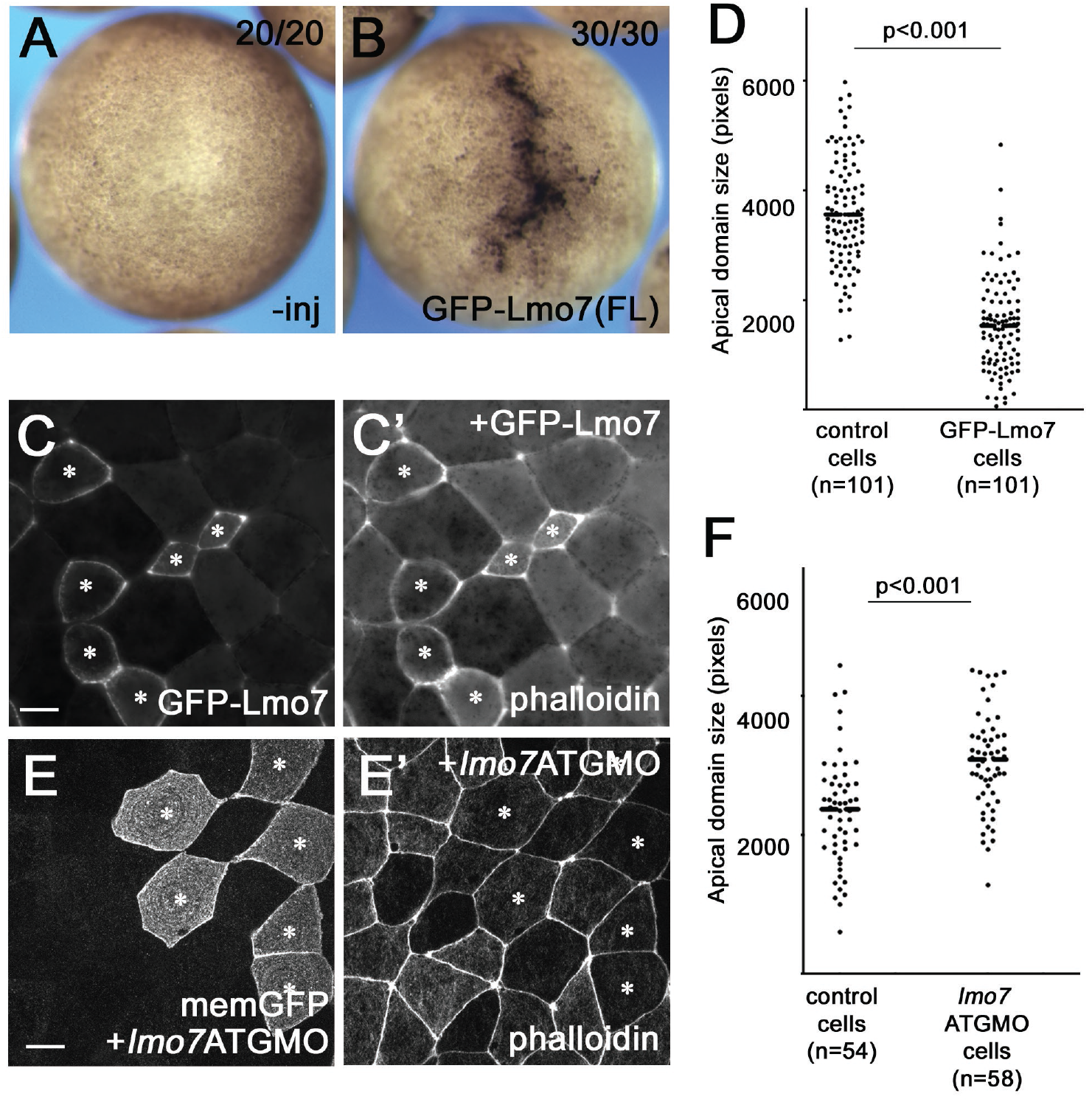
Lmo7 induces apical constriction in *Xenopus* ectoderm. (A, B) GFP-Lmo7 mRNA was injected into two animal blastomeres of 4-8 cell stage embryos. (A) Control uninjected embryo, stage 11. (B) Injection of GFP-Lmo7 mRNA (1 ng) triggers apical pigment granule accumulation. (C, C’), Reduced apical domain size in cells expressing GFP-Lmo7 (200 pg mRNA). GFP fluorescence and phalloidin staining are shown for ectoderm of stage 11 embryo. Asterisks show GFP-Lmo7-expressing cells. (D) Quantification of apical domain size in cells expressing GFP- Lmo7 (n=101) and adjacent control cells (n=101). Images were taken from at least 5 embryos. (E, F) *lmo7* knockdown increases apical domain size. *Lmo7* ATGMO (30 ng) was injected into one blastomere of 4-cell stage embryos together with membrane-tethered GFP (memGFP) mRNA (100 pg). Embryos were co-stained with phalloidin at St 11. Asterisks show *lmo7*MO-containing cells. (F) Quantification of apical domain size in cells with *lmo7* ATGMO (n=58) and control (-GFP) cells (n=54) from more than five different embryos. Statistical significance of the difference between the median values was assessed by the Student’s t-test. Scale bars: 10 μm in A, C E and G. 2 μm in B, D and F.

Phalloidin staining and the quantification of apical domain size confirmed the apical constriction in individual ectoderm cells (Fig. 2C, C’, D). To deplete Lmo7, a translation-blocking morpholino oligonucleotide (*lmo7-*ATGMO) was injected in the ectoderm. Although no overt effect on ectoderm pigmentation was apparent, *lmo7* knockdown expanded the apical domain (Fig. 2E, E’, F). This effect of *lmo7*MO was rescued by co-injection of GFP-Lmo7 mRNA (Fig. S1). This is consistent with the Lmo7 function in apical constriction.

### Lmo7 reorganizes the actomyosin network to regulate NMII-dependent contractility at apical junctions

Highly organized actomyosin network at AJCs is known to generate the contractile force during apical constriction. The Rho-ROCK-NMII signaling module plays a key role in this process (Martin and Goldstein, 2014). We examined actomyosin bundles at the borders of Lmo7-expressing cells and compared them with those between non-expressing cells (Fig. 3A-L). Lmo7-expressing cells exhibited thick double bands of Flag-Lmo7 in a sarcomere-like pattern (Fig. 3C, G, K). Flag-Lmo7 induced the formation of intense F-actin double bands that largely overlapped with Lmo7 (Fig. 3B-C”). The effect was less pronounced in embryos injected with lower dose of *lmo7* mRNA (Fig. 1D, F), suggesting dose-dependent effects. The appearance of thick F-actin bundles was accompanied by accumulation of phosphorylated myosin regulatory light chain (pMRLC) (Fig. 3E-H), NMIIA (Fig. 3I-L) and exogenously expressed *α*-actinin 4 (Fig. S2). These results suggest that Lmo7 promotes actomyosin bundle formation at apical junctions.

**Figure 3.**
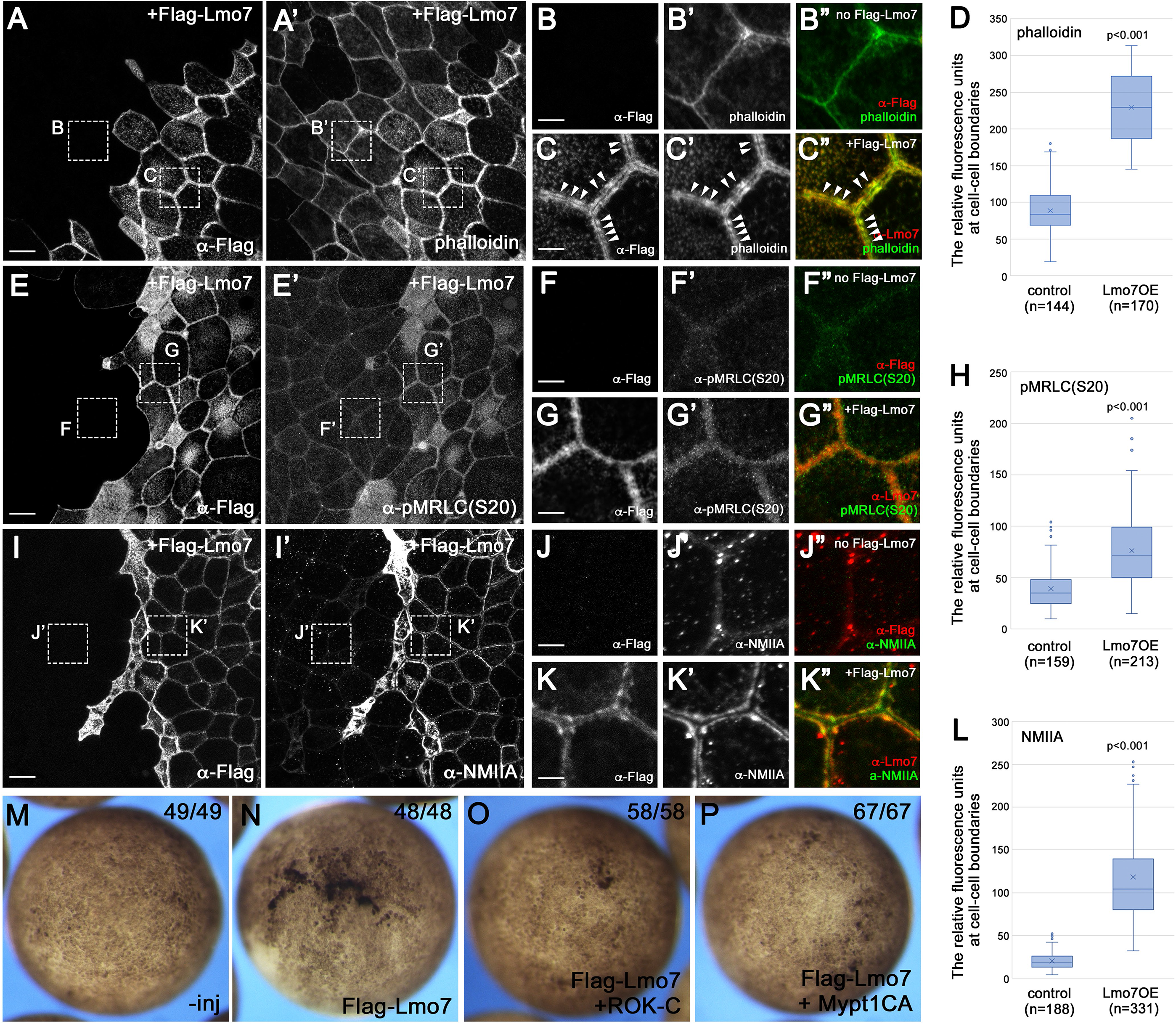
Lmo7 promotes actomyosin bundle formation at apical junctions through the Rho-ROCK-MyoII pathway. (A-L) Effects of Flag-Lmo7 overexpression on actomyosin networks. Flag-Lmo7 mRNA (1 ng) was injected into one blastomere of 4-8 cell stage embryos. Embryos were co-stained with phalloidin, anti- Flag, pMRLC or NMIIA antibodies as indicated. The superficial ectoderm was imaged at stage 11. (A- C”) Flag-Lmo7 induces the formation of thick and dense F-actin bundles. Areas marked by rectangles in A-A’ are enlarged in B-C”. (C-C”) Lmo7 double bands formed between two adjacent Flag-Lmo7 cells largely overlap with F-actin double bands. Note periodic sarcomere-like structures (arrowheads). (D) Fluorescence intensity of phalloidin was measured at 3-10 locations at a border between two adjacent control cells (control) or two Lmo7 expressing cells (Lmo7OE). (E-G”) MRLC phosphorylation at Ser 20. Areas marked by rectangles in E-E’ are enlarged in F-G”. (H) Fluorescence intensity of pMRLC(S20) staining related to images in E-G” (I-K”) NMIIA accumulation. Areas marked by rectangles are enlarged in J-K”. (L) Fluorescence intensity of NMIIA staining related to images in I-K”. (M-P) Rho-ROCK-NMII signaling is necessary for Lmo7-induced apical constriction. (M) Control uninjected embryo, (N) Flag- Lmo7 mRNA (500 pg)-injected embryos. (O) Coinjection of Flag-Lmo7 mRNA (500 pg) and ROK-C mRNA (200 pg) (P) Coinjection of Flag-Lmo7 mRNA (500 pg) and Mypt1CA mRNA (50 pg). Images are representative of three experiments. Scale bars: 10 μm in A, A’, E, E’, I and I’. 2 μm in B-C”, F-G” and J-K”.

We next asked whether Lmo7-mediated apical constriction involves NMII, a known driver of early morphogenetic processes in *Xenopus* (Skoglund et al., 2008). We inhibited MRLC phosphorylation by co-expression of RNAs encoding a dominant-interfering form of ROCK (ROK-C) or the constitutively active subunit of myosin phosphatase Mypt1 (Mypt1CA). Both have been shown to decrease MRLC phosphorylation that is necessary for NMII bipolar filament formation (Feng et al., 1999; Marlow et al., 2002; Weiser et al., 2009). We found that co-expression of either RNA suppressed Lmo7-mediated pigment granule accumulation in *Xenopus* ectoderm (Fig. 3M-P). These results suggest that Rho-ROCK-NMII signaling is required for Lmo7-dependent apical constriction.

### Structure-function analysis of Lmo7-dependent apical constriction

Lmo7 has the calponin homology domain (CH), the <u>d</u>omain of <u>u</u>nknown function DUF4757, the *α*-actinin-binding region, the PDZ domain, two coiled-coil domains and the LIM domain which binds to Afadin (Ooshio et al., 2004) (Fig. 4A). To define the domains of Lmo7 required for apical constriction, various Lmo7 constructs were expressed in the superficial ectoderm (Fig. S3)(Fig. 4B-I). We found that the removal of the N-terminal CH domain (Fig. 4B), the C-terminal LIM domain (Fig. 4F), or the coiled-coil domain and the PDZ domain (Fig. 4C) did not affect the apical constriction. On the other hand, the *α*- actinin binding region was necessary for this activity (Fig. 4D). DUF4757 was also necessary but not sufficient on its own (Fig. 4, G, I, K, K’, L). Lmo7(aa 242-709) containing both DUF4757 and the *α*- actinin-binding region was the shortest Lmo7 fragment that triggered apical constriction (Fig. 4H) and reduced apical domain size in individual cells (Fig. 4J, J’, L). Taken together, these studies demonstrate essential roles of DUF4757 and the *α*-actinin-binding region in Lmo7-mediated apical constriction.

**Figure 4.**
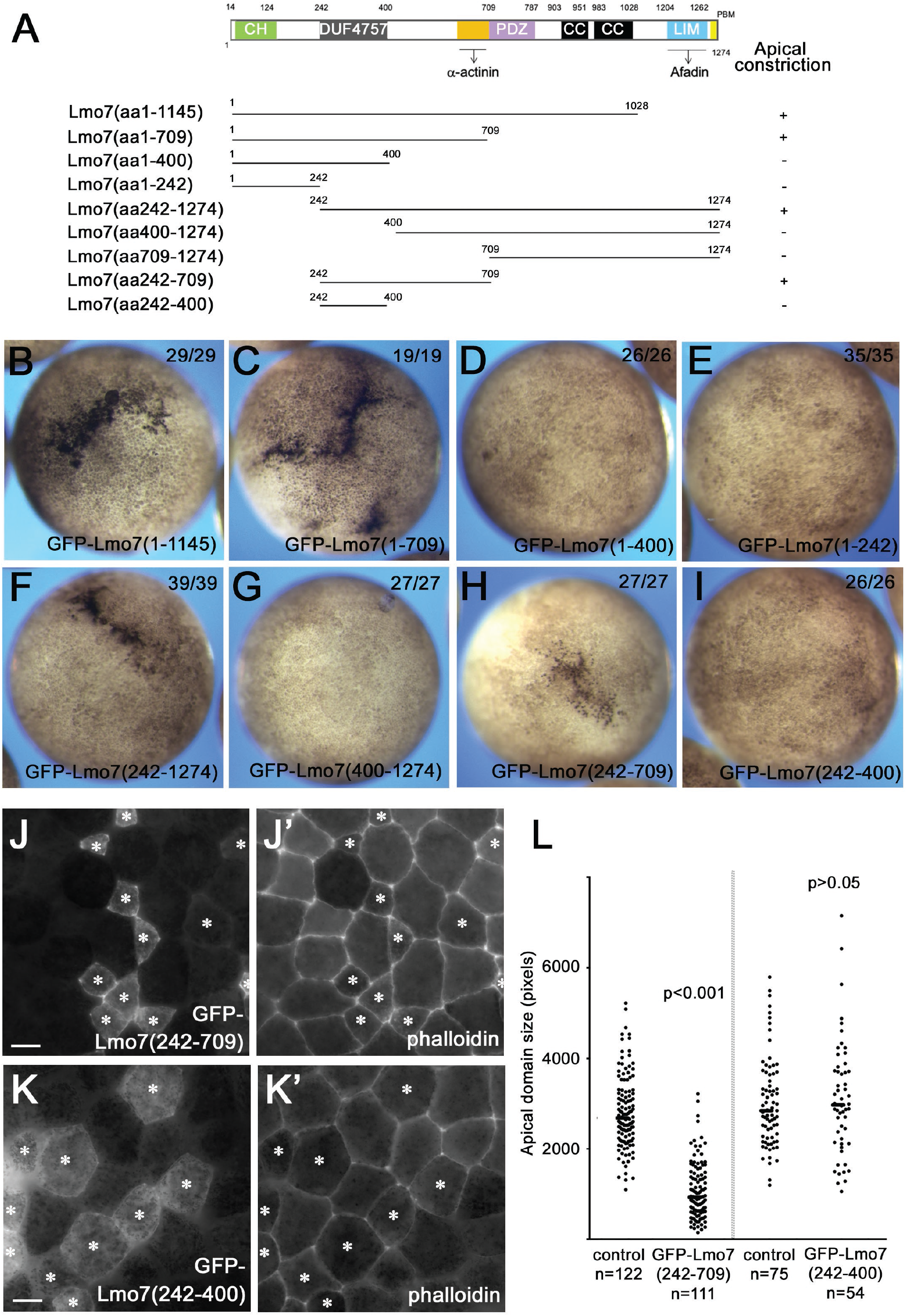
Identification of Lmo7 domains required for apical constriction. (A) Domain structure of Lmo7 and the mutants used in this study. Lmo7 has the N-terminal Calponin homology (CH) domain, the DUF4757 domain, the *α*-actinin binding region, the PDZ domain, two coiled-coil domains and the C-terminal LIM domain. (B-I) Representative images of stage 11 embryos injected with mRNA encoding various Lmo7 constructs (1 ng). Apical constriction was assessed by pigment granule accumulation. Apical constriction induced by Lmo7 constructs lacking the LIM domain (B), coiled-coil and PDZ domain (C). Deletion of the *α*-actinin-binding region (D) and DUF4757 (E) eliminates the activity, whereas CH domain deletion has no visible effect (F). Lmo7(aa 400-1274) lacking DUF4757 has no activity (G). The region containing the DUF4757 domain and *α*-actinin binding region Lmo7(aa 242-709) can induce pigment granule accumulation (H), but the DUF4757 domain has no activity on its own (I). (J-L) GFP-Lmo7(aa 242-709) but not GFP-Lmo7(aa 242-400) reduces apical domain size. GFP-Lmo7 deletion mutants were mosaically expressed by mRNA injection (200 pg, asterisks) in 4-8-cell stage embryos. GFP fluorescence and phalloidin staining are shown in stage 11 ectoderm. (L) Quantification of apical domain size for experiments shown in J-K’. Apical domain size of control cells was measured only in cells that shared at least one cell boundary with cells expressing GFP- Lmo7 mutants. Images for scoring were taken from at least five embryos. Data are representative of more than three independent experiments. Scale bars: 10 μm in J and K.

### Lmo7 binds and recruits NMII to apical junctions via DUF4757

To gain further insight into potential molecular mechanism of Lmo7-dependent apical constriction, we examined whether Lmo7 physically interacts with NMII in HEK293T cells. We found both endogenous NMIIA and NMIIB in Flag-Lmo7 pull-downs (Fig. 5A, B). DUF4757 was both required and sufficient for this binding, suggesting that Lmo7 effect on the circumferential actomyosin belt may be due to this DUF4757 interaction with NMII heavy chains. Alanine substitution of the conserved WQ-WK amino acids in DUF4757 (Fig. 5A) eliminated the ability of Lmo7 to recruit NMIIA to cell junctions (Fig. 6A- C’, G, H). These results indicate that DUF4757 is required for NMII accumulation at apical junctions. Importantly, GFP-Lmo7(AAAA) and GFP-Lmo7(AAWK) did not induce apical constriction in ectodermal cells (Fig. S4). By contrast, Lmo7(aa 242-709) triggered apical constriction and recruited NMIIA to apical junctions (Fig. 6D-F’, I), although less efficiently than full-length Lmo7 (compare Fig. 6E-E’, G and F-F’, I). Notably, Lmo7(aa 242-709) did not increase F-actin accumulation at apical junction (Fig. S5). These observations suggest that Lmo7 induces apical constriction by recruiting NMII in the proximity of apical junctions via its NMII-binding domain.

**Figure 5.**
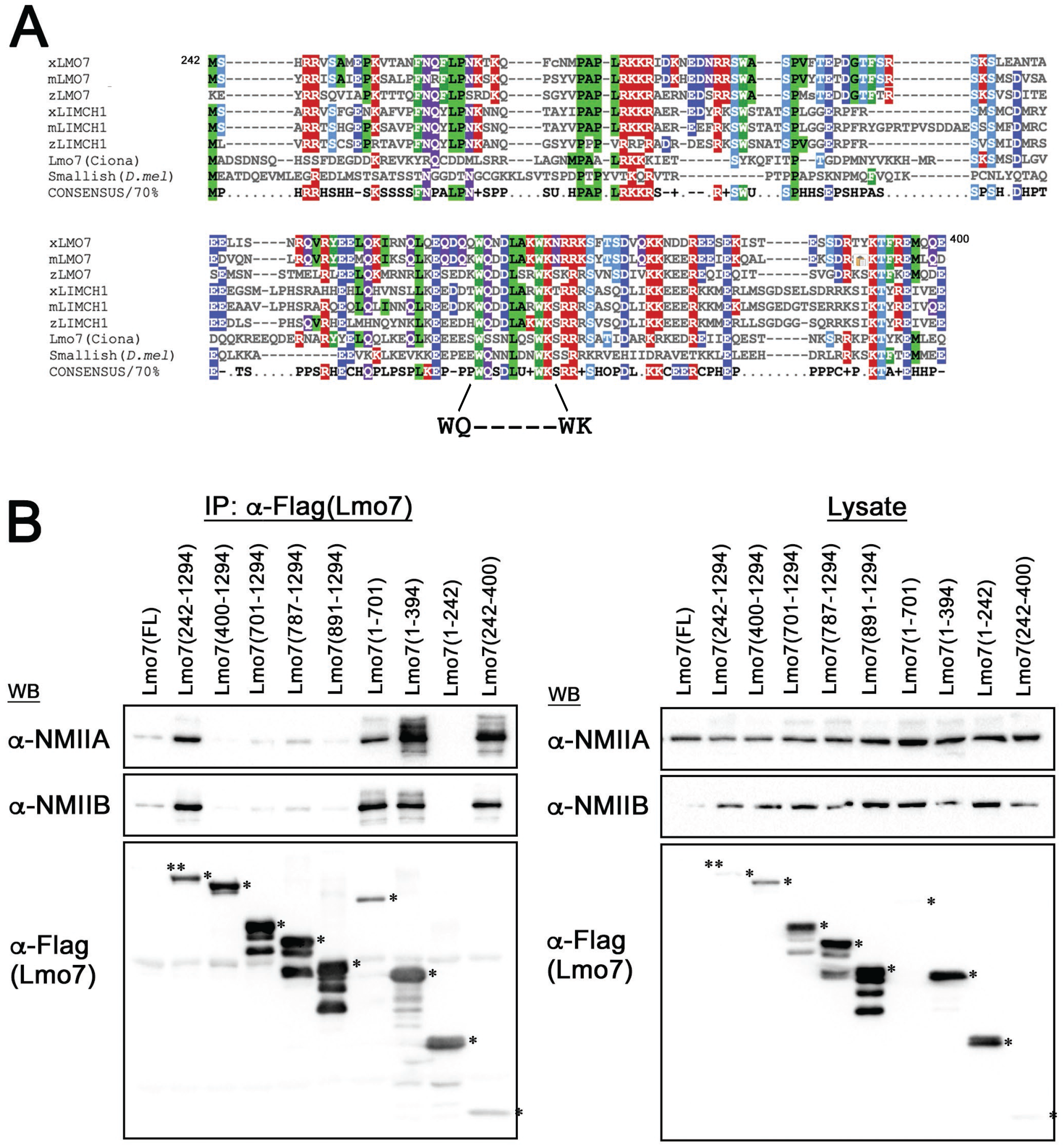
The DUF4757 domain binds to NMII heavy chain. (A) The alignment of DUF4757 sequences. *Xenopus laevis* Lmo7 (NM_001135230), *Mus musculus* Lmo7 (XM_006519190), *Danio rerio* Lmo7 (XM_021478660), *Xenopus laevis* LIMCH1 (NM_001096206), *Mus musculus* LIMCH1 (NM_001001980), *Danio rerio* LIMCH1 (XM_009291008), *Ciona intestinalis* Lmo7 (XM_002128910) and *Drosophila melanogaster* Smallish (NM_169014). Conserved positively charged, negatively charged, hydrophobic, or hydrophilic amino acids are marked by red, blue, green and purple, respectively. Ser or Thr residues are marked by light blue. Consensus sequence based on amino acid similarity (>70%) is shown at the bottom. The conserved WQ-WK sequence used for AA-AA substitution is highlighted. (B) NMIIA and NMIIB coprecipitate with Flag- Lmo7 from transfected HEK293T cell lysates. Asterisks indicate protein bands corresponding to full- length or deletion mutant forms of Lmo7. Note that Lmo7(aa 242-400) containing DUF4757 is sufficient to coprecipitate endogenous NMIIA and NMIIB.

**Figure 6.**
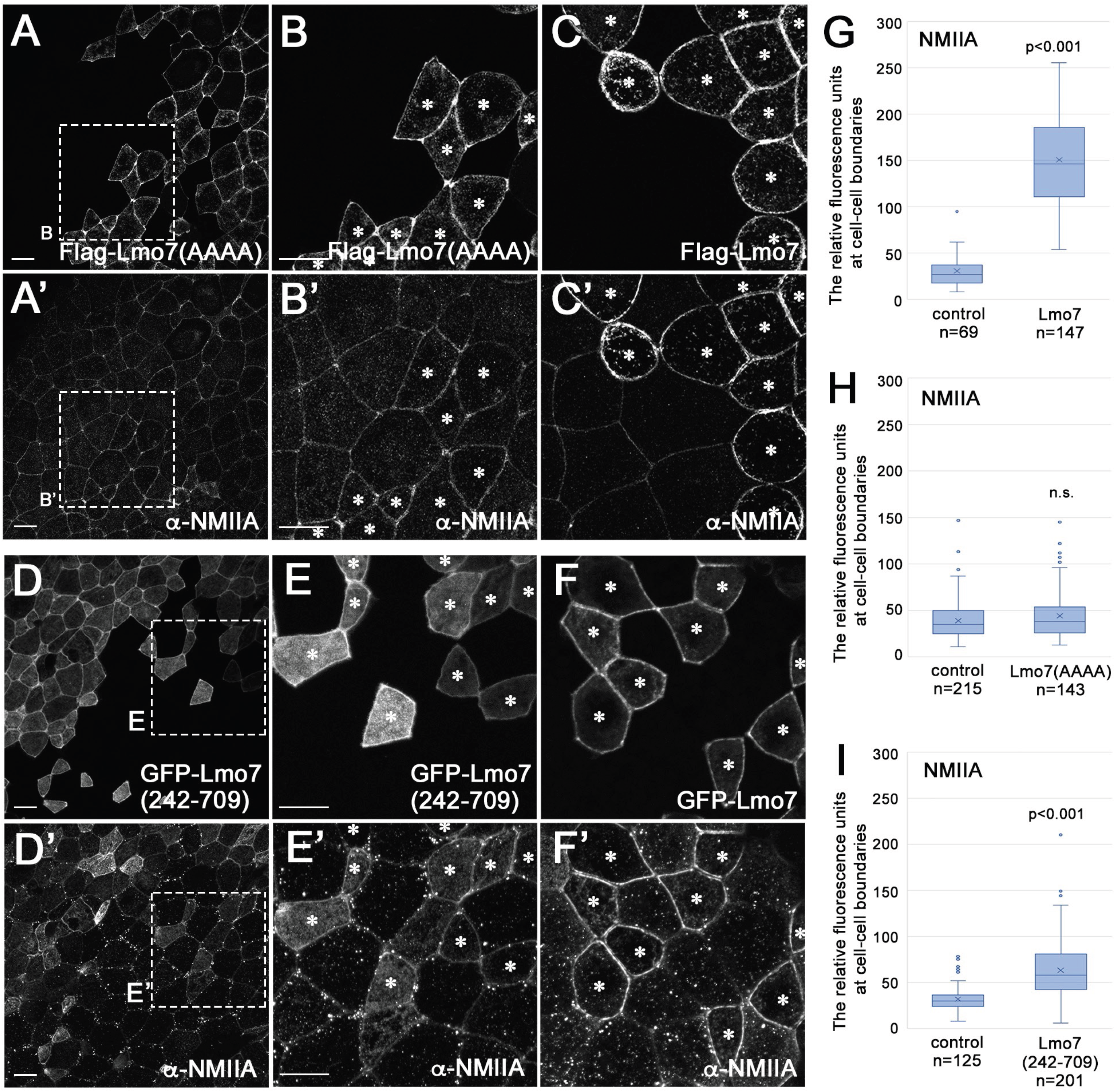
Lmo7 recruits NMII to apical junctions through the DUF4757 domain. Embryos were injected with RNAs (0.5-1 ng) encoding full-length or mutant forms of Lmo7 and cultured until stage 11. Superficial ectoderm stained with anti-Flag, -GFP or -NMIIA antibodies is shown. Cells expressing Flag-Lmo7 constructs are marked by asterisks. (A-B’) Lmo7(AAAA) does not cause NMIIA enrichment at apical junctions. Areas marked by rectangles in A and A’ are enlarged in B and B’. (C-C’) Comparable images of embryo expressing Flag-Lmo7. (D-E’) Lmo7(aa 242-709) increases NMIIA accumulation. Areas marked by rectangle in D and D’ are enlarged in E and E’. (F, F’) Comparable images of embryo expressing GFP-Lmo7. (G-I) Quantification of NMIIA accumulation at apical junctions in cells expressing GFP-Lmo7 (G), GFP-Lmo7(AAAA)(H), or GFP-Lmo7(aa 242-709)(I). Fluorescence intensity of NMIIA was measured at 3-10 locations within individual cell-cell boundaries. Note that GFP-Lmo7(242-709) recruits NMIIA less effectively than full-length Lmo7. Scale bars: 10 μm.

The role of Lmo7 in NMII recruitment to apical junctions has been confirmed in loss-of-function experiments. We observed reduction of endogenous NMIIA in the cells depleted of Lmo7 (Fig. 7B-C’). Nevertheless, these cells preserved normal morphology (Fig. 7C, C’) and the junctional distribution of ZO-1 (Fig. 7D-E’). Taken together, our findings suggest that Lmo7 functions to maintain NMII at apical junctions.

**Figure 7.**
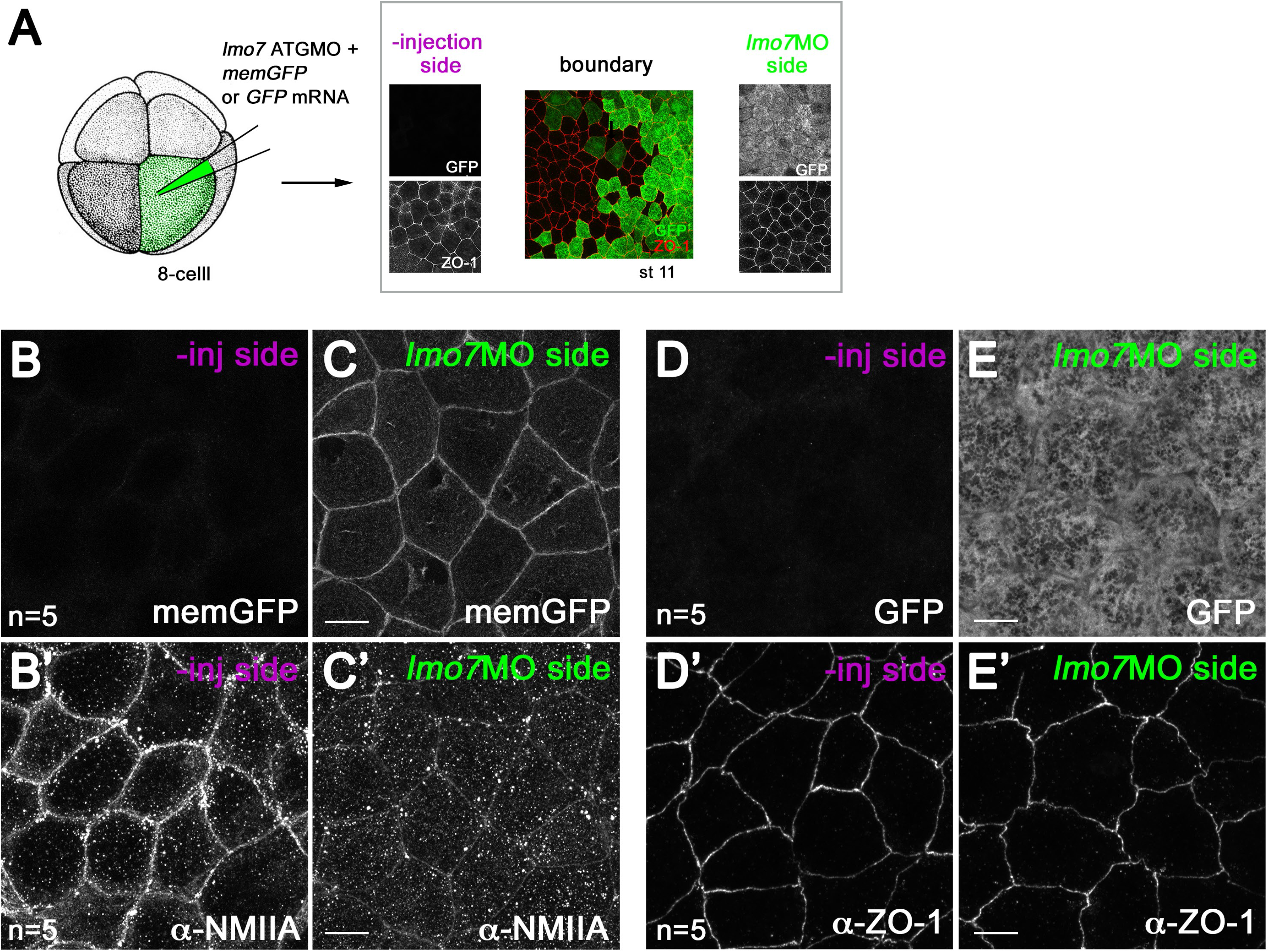
Lmo7 knockdown reduces NMIIA accumulation at apical junctions. (A) Schematic diagram of the experimental design. *Lmo7* ATG MO (30 ng) was co-injected into one ventral blastomere of 4-8 cell stage embryos together with mRNA encoding GFP or membrane-tethered GFP (memGFP)(100 pg) for lineage tracing. Stage 11 embryos were stained with anti-NMIIA or ZO-1 antibodies. (B-C’) NMIIA enrichment at apical junctions is reduced in *lmo7*MO cells. Note that *lmo7*MO cells maintain normal cell morphology labeled by memGFP (C, C’). (D-E’) No significant change in ZO- 1 distribution at TJs in *lmo7*MO cells. Scale bars: 10 μm.

### Lmo7 induces tension-dependent relocalization of Wtip at apical junctions

By recruiting NMII to actomyosin bundles, Lmo7 may generate mechanical force at apical junctions. We took advantage of the observations that cytoplasmic and cortical aggregates of Ajuba family proteins, such as Wilms-tumor-1-interacting protein (Wtip), dissociate in response to cytoskeletal tension (Chu et al., 2018; Rauskolb et al., 2014). We used this localization change as a readout of increased mechanical tension at apical junctions. To test whether Lmo7 can generate force, we injected Flag-Lmo7 mRNA into a subpopulation of RFP-HA-Wtip-expressing cells (Fig. 8A). RFP-HA-Wtip puncta were observed in the cytoplasm and cell junctions of majority of control cells (Fig. 8B-C”, E). These puncta were eliminated by Lmo7 (Fig. 8B-B”, D-D”, E). Whereas the physiological significance of Wtip puncta formation requires further investigation, these results suggest that Lmo7 increases tension at apical junctions.

**Figure 8.**
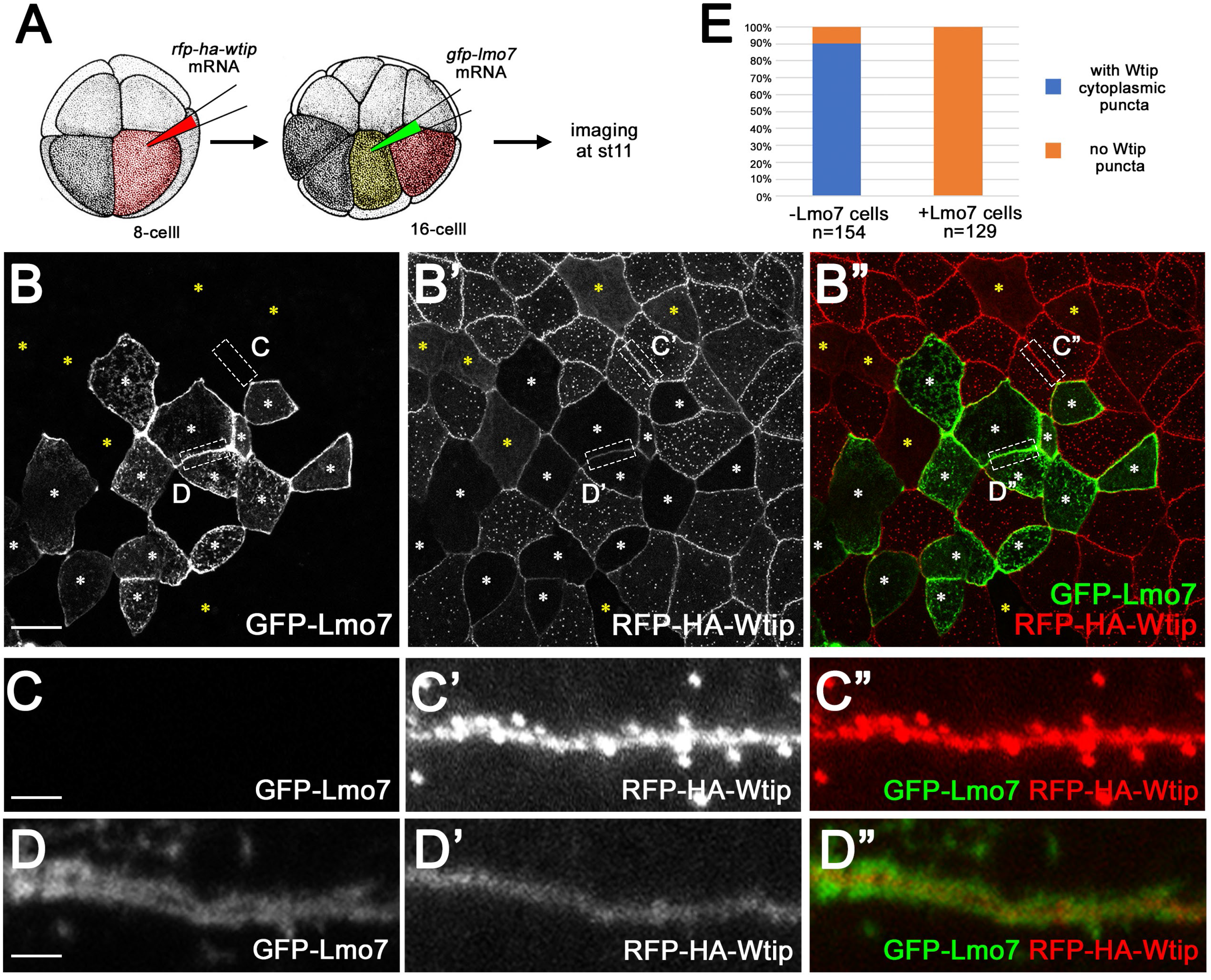
Lmo7 induces the tension-based localization change of RFP-HA-Wtip at apical junctions. (A) Schematic diagram of the experimental design. mRNA encoding RFP-HA-Wtip (200 pg) was injected into one ventral blastomere of 4-8 cell stage embryos (in red). At around 16-32 cell stage, mRNA encoding GFP-Lmo7 (500 pg) was injected into one of the blastomeres with RFP-HA-Wtip (in yellow). This protocol results in approximately 25-50 % cells coexpressing RFP-HA-Wtip and GFP-Lmo7 at atage 11. (B-D”) Lmo7 co-expression eliminates RFP-HA-Wtip puncta both in the cytoplasm and near apical junctions. White asterisks show cells expressing both GFP-Lmo7 and RFP-HA-Wtip. Note that approximately 10 % of cell population only expressing Wtip-HA-RFP does not have Wtip-HA-RFP puncta in the cytoplasm (marked by yellow asterisks). Areas of rectangles in B-B” are enlarged in C-D”. (C-C”) RFP-HA-Wtip puncta are present near apical junctions in control cells. (D-D”) RFP-HA-Wtip puncta are absent near apical cell-cell junctions in GFP-Lmo7 cells. Note that RFP-HA-Wtip forms a smooth line between the Flag-Lmo7 double bands. (E) Quantification of RFP-HA-Wtip puncta formation near apical junctions in the presence or absence of GFP-Lmo7. Scale bars: 10 μm in B. 2 μm in C-D.

## Discussion

In this study, we identify Lmo7 as a novel regulator of apical constriction in *Xenopus* embryonic ectoderm. Lmo7 localizes to peripheral actomyosin cables that undercoat apical junctions. Embryonic cells expressing Lmo7 form thicker and denser actomyosin bundles as compared to control ectoderm. We found that Lmo7 binds NMII heavy chains via a conserved domain, DUF4757. Analysis of Lmo7 deletion mutants revealed a correlation between the apical constriction inducing activity and the recruitment of NMII to apical junctions. We propose that Lmo7 triggers apical constriction by promoting NMII incorporation into perijunctional actomyosin networks.

The striking localization of Lmo7 to two cortical bands on both sides of the apical cell-cell junction is similar to the distribution of NMIIA in the normal tissue. The double bands of NMII have been described for non-sensory epithelial cells of the mammalian inner ear (Ebrahim et al., 2013). Similar organization of NMII was induced in MDCK cells by depletion of ZO proteins, suggesting crosstalk between apical junctions and actomyosin contractility (Choi et al., 2016; Fanning et al., 2012; Tokuda et al., 2014; Van Itallie et al., 2009). Lmo7 is a scaffold protein that can mediate such crosstalk. In gain-of-function experiments, Lmo7 promoted the formation of organized perijunctional actomyosin bundles, culminating in apical constriction. Our knockdown experiments revealed increased apical domain size and reduced NMIIA at apical junctions of ectoderm cells. Together, these observations lead us to propose that vertebrate Lmo7 is responsible for the organization and contractility of the circumferential actomyosin belt. Consistent with this hypothesis, *Drosophila* embryos lacking Smallish, the only fly orthologue of Lmo7, are defective in epithelial morphogenesis (Beati et al., 2018). By contrast, *lmo7* knockout mice exhibit limited developmental defects in specific epithelia (Du et al., 2019; Lao et al., 2015; Tanaka-Okamoto et al., 2009). These findings indicate that the loss of Lmo7 is functionally compensated by other genes, such as its paralogue Limch1 (Lin et al., 2017). Double knockout of *lmo7* and *limch1* will be necessary to properly evaluate their loss-of-function phenotypes in vertebrates.

Our experiments show that Lmo7 triggers apical constriction in *Xenopus* ectoderm and this activity requires the recruitment of NMII heavy chains via DUF4757, a domain with previously unknown function. These findings are consistent with an earlier study showing that Limch1 associates with NMII and promotes stress fiber formation in Hela cells (Lin et al., 2017). Of note, DUF4757 is disrupted in certain splicing variants of Lmo7 (data not shown) and only a portion of it is present in Smallish. In addition to recruiting NMII to apical junctions, Lmo7 appears to act as a scaffold to position F-actin, NMII and *α*-actinin at actomyosin bundles and organize them in a sarcomere-like pattern. Among the domains that may interact with F-actin and contribute to the ability of Lmo7 to modulate actomyosin cables are the CH domain and the Afadin-binding LIM domain (Ooshio et al., 2004). Surprisingly, Lmo7(aa 242-709), lacking these domains, triggered apical constriction but did not promote F-actin accumulation as compared to the full-length protein. Further analysis is needed to gain further insights into the scaffolding functions of Lmo7 domains in actomyosin network regulation.

The molecular mechanism of Lmo7-dependent apical constriction requires further investigation. Lmo7 may stimulate NMII head domain ATPase activity to increase contractility, modulate NMII heavy chain phosphorylation (Conti et al., 1991; Dulyaninova et al., 2007; Even-Faitelson and Ravid, 2006; Murakami et al., 1998) or NMII interaction with other proteins (Dahan et al., 2012; Dulyaninova et al., 2005; Elliott et al., 2012; Garrett et al., 2006). The effect of Lmo7 on actomyosin bundles may be relevant to mechanotransduction processes at apical junctions. Lmo7 has been reported to respond to the mechanical force (Hashimoto et al., 2019). Lmo7 binds Afadin that was implicated in force transmission at cell-cell junctions (Choi et al., 2016; Manning et al., 2019; Yu and Zallen, 2020). Supporting this hypothesis, we found that Lmo7 influences the distribution of Wtip, a sensor of mechanical tension (Chu et al., 2018).

The remaining challenge is to determine how the interplay of Lmo7, Afadin and other junctional proteins leads to effective generation, sensing and transmission of mechanical forces during epithelial morphogenesis.

## Materials and Methods

### Plasmids

*Xenopus laevis* Lmo7 cDNA clone was obtained from Dharmacon. Full-length and truncated forms of Lmo7 were PCR amplified and cloned into a pCS107-Flag, -HA or -GFP expression vector using standard protocols. Primers used for the amplification are in Supplemental Table 1. pCS2-RFP-HA-Wtip was previously described (Chu et al., 2018). pCS2-GFP-NMIIA and pCS2-GFP-NMIIB were subcloned from CMV-GFP-NMHC II-A, CMV-GFP-NMHC II-B (Addgene plasmids # 11347, 11348)(Wei and Adelstein, 2000). pCS2-GFP-*α*-actinin 4 was constructed from pCS2-hiActTS-GR (a gift of Tatsuo Michiue). pCS2-ROK-C and pCS2-Mypt1CA were from Florence Marlow (Marlow et al., 2002) and David Kimelman (Weiser et al., 2009), respectively.

### *Xenopus* embryo culture and microinjections of RNA and morpholinos

Wild-type *Xenopus laevis* adults were purchased from Nasco, maintained and handled according to the recommendations in the Guide for the Care and Use of Laboratory Animals of the National Institutes of Health. A protocol for animal use was approved by the Institutional Animal Care and Use Committee (IACUC) at Icahn School of Medicine at Mount Sinai. *In vitro* fertilization and embryo culture were performed as described previously (Dollar et al., 2005). Embryo staging was determined according to Nieuwkoop and Faber (1967). For microinjections, embryos were transferred into 3 % Ficoll 400 (Pharmacia) in 0.5x Marc’s modified Ringer’s (MMR) solution (50 mM NaCl, 1 mM KCl, 1 mM CaCl2, 0.5 mM MgCl2 and 2.5 mM HEPES (pH 7.4))(Peng, 1991). Linearized plasmid DNA was used to synthesize mRNA using mMessage mMachine SP6 Transcription kit (Invitrogen). Synthesized mRNA was purified by LiCl precipitation. mRNA in 5-10 nl of RNase-free water (Invitrogen) was microinjected into one or two blastomeres of four to sixteen cell stage embryos. Morpholino oligonucleotide (MO) specific for Lmo7 was purchased from Gene-Tools, and had the following sequence: 5’- GAATTTTCATTCCATTCCATTG-3’. GFP or memGFP mRNA was coinjected with MO for lineage tracing. Imaging of GFP-negative cells was done at least 20 cells away from the boundary region, considering the difference in diffusion of RNA and MO.

### Cell culture and transfection

HEK293T were purchased from ATCC and maintained in Dulbecco’s modified eagle media (DMEM) (Corning) supplemented with 10 % fetal bovine serum (Sigma) in a 37 C incubator with 5% CO2.

HEK293T cells grown in 60 mm dish to 70-80 % density were transfected with a mixture of 4 μg plasmid DNA and 20 μg polyethylenimine (Polysciences). Cell lysates were extracted 24 hrs after transfection.

### Immunoprecipitation and immunoblotting

Cell lysates were extracted from HEK293T cells using RIPA buffer (50 mM Tris-HCl pH 7.4, 150 mM NaCl, 1 % NP-40, 0.5% sodium deoxycholate, 0.1 % SDS) supplemented with Protease Inhibitor Cocktail III (Calbiochem) and PhosStop phosphatase inhibitor cocktail (Roche). Cell lysates were incubated with anti-Flag M2 agarose (Sigma) at 4°C at least 6 hours. Agarose beads were washed three times in Tris-buffered saline containing 0.05 % Triton X-100 (TBST). Agarose beads were heated at 95°C for 5 min in SDS sample buffer containing 5 % *β*-mercaptoethanol (Sigma). After SDS-PAGE and transfer to a nitrocellulose membrane with a 0.2 μm pore size (RioRad), immunoblotting was done with the following antibodies: rabbit anti-NMIIA pAb (BioLegend, #POLY19098, 1:500), rabbit anti-NMIIB pAb (BioLegend, #POLY19099, 1:500), mouse anti-DYKDDDDK mAb clone 2H8 (Cosmo Bio USA, #KAL- K0602, 1:1000) and mouse anti-GFP mAb clone B2 (Santa Cruz Biotechnology, #sc-9996, 1:500). HRP-conjugated secondary antibodies against mouse or rabbit IgG were from Cell Signaling Technology (#7074, #7076, 1:5000). Chemiluminescent signals were acquired using Clarity ECL Western Blotting Substrates (Bio-Rad) on the ChemiDoc MP Imaging System (Bio-Rad).

*Xenopus* embryo lysates were made by incubating 20 embryos in 400 μl lysis buffer (50 mM Tris-HCl (pH 7.6), 50 mM NaCl, 1 mM EDTA, 1 % Triton X-100) supplemented with protease inhibitor and protein phosphatase inhibitor cocktails as described above. After centrifugation for 3 min at 16,000 g, the supernatant was mixed with SDS-sample buffer, followed by standard SDS-PAGE and western blot protocol. Experiments were repeated at least three times.

### Examination of apical constriction

RNAs encoding Lmo7, ROK-C or Mypt1-CA were injected into two animal-ventral blastomeres of 4-cell embryos. For analysis of pigment granule accumulation, embryo images were captured using a Leica stereo microscope equipped with a color CCD camera. Each experiment included 20-30 embryos per condition. Experiments were repeated at least three times.

For measurement of apical domain size, cells expressing GFP-Lmo7 or its deletion mutant forms and their surrounding cells were outlined manually using a free-hand line tool based on phalloidin staining. ImageJ was used to quantify the apical domain size. Each experiment included 10-20 embryos per condition. Experiments were repeated at least twice.

### Wholemount immunocytochemistry

For formaldehyde fixation, *Xenopus* embryos were devitellinized manually using forceps and fixed in MEMFA (0.1 M MOPS (pH 7.4), 2 mM EGTA, 1 mM MgSO4, and 3.7% formaldehyde)(Harland, 1991) for 60 min at room temperature. After washing in phosphate buffered saline (PBS), embryos were permeabilized in 0.1 % Triton in PBS for 30 min. For trichloroacetic acid (TCA) fixation, devitellinized embryos were fixed in ice-cold 10% TCA for 30 min. Embryos were then incubated in 1 % BSA in TBS at 4°C overnight for blocking. After that, embryos were incubated with primary antibodies in 1 % BSA, 1 % DMSO in PBS or TBS at 4C overnight. Antibodies used in this study are the following: rabbit anti-

NMIIA pAb (BioLegend, POLY19098, 1:300), mouse anti-DYKDDDDK mAb clone 2H8 (Cosmo Bio USA, KAL-K0602, 1:500), rat anti-HA mAb clone 3F10 (Roche, 118673230002, 1:100), and chicken anti-GFP pAb (abcam, ab13970, 1:500). After wash in PBS or TBS three times, embryos were incubated with secondary antibodies conjugated with either Alexa 488, Cy3 or Cy2 fluorescent dyes in 1% BSA, 1% DMSO in PBS at 4C overnight. Alexa 488, Cy3, or Cy2-conjugated secondary antibodies against mouse, rat and rabbit IgG or chicken IgY are obtained from Invitrogen (A-32723, A-11034, 1:300) or Jackson ImmunoResearch Laboratory (711-165-152, 715-165-151, 712-165-153, 703-225-155, 1:500). AlexaFluor 555-phalloidin (Invitrogen, A34055) was added to secondary antibody solution to label F-actin. After wash in PBS or TBS three times, embryos are transferred in 25 % glycerol in PBS. The animal pole side of embryos was dissected manually by forceps and mounted on slide glasses using Vectashield mounting medium (Vector). Imaging was done using Zeiss AxioImager upright microscope with Zeiss Axiocam CCD camera or Zeiss LSM880 confocal microscope system. Images were captured from at least 5 independent embryos. Experiments were repeated at least three times. Captured images were processed and quantified using ImageJ software.

### Quantification and statistical analyses

Quantification of fluorescence signals was performed by analyzing individual single plane images. Integrated fluorescence intensity of immunostaining was measured using an ImageJ plugin. The Student’s t-test was used to analyze statistical significance of the difference between the samples.

## Acknowledgements

We thank Robert Adelstein, Florence Marlow, Tatsuo Michiue and David Kimelman for plasmids, Pamela Mancini for help with Lmo7 mutagenesis at the early stages of this project and members of the Sokol laboratory for discussions. We acknowledge the help from the ISMMS Microscopy Core facility. This research was supported by the NIH grant R35GM122492 to SYS.

## Data availability

n.a.

