## Supplemental information for "Lmo7 recruits myosin II heavy chain to induce apical constriction in *Xenopus* ectoderm"

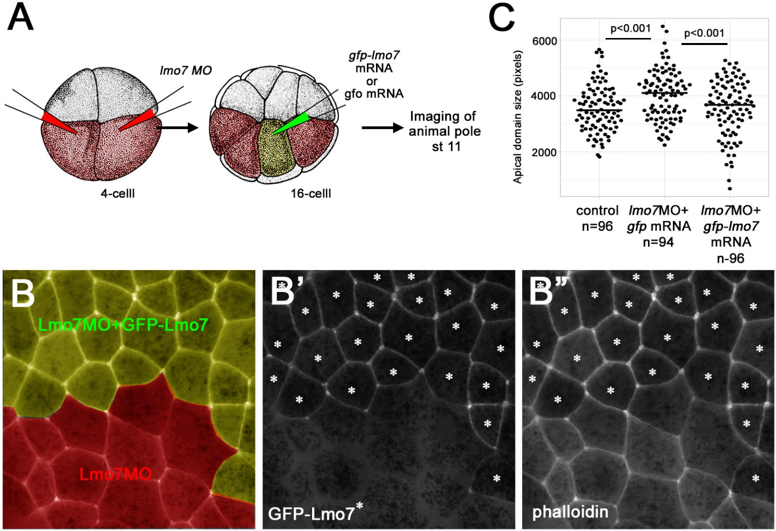
A
lmo7 MO
gfp-lmo7
mRNA
or
gfo mRNA
Imaging of
animal pole
st 11
4-celll
16-celll
C
p<0.001
p<0.001
6000
4000
2000
Apical domain size (pixels)
control
n=96
lmo7MO+
gfp mRNA
n=94
lmo7MO+
gfp-lmo7
mRNA
n-96
B
Lmo7MO+GFP-Lmo7
Lmo7MO
B'
GFP-Lmo7*
B''
phalloidin

### Figure S1

**Supplemental Figure 1. GFP-Lmo7 co-expression rescues increased apical domain expansion in *lmo7* morphants**

(A) Schematic diagram of the experimental design. *lmo7* MO (30 ng) was injected into both ventral blastomeres of four-cell embryos (red). At 16-32 cell stage, mRNA encoding control GFP (100 pg) or GFP-Lmo7 (100 pg) was injected into one of the ventral blastomeres (yellow). The sequential injection allows to compare MO-injected cells (the majority of animal hemisphere cells) with the cells double-injected with MO and GFP or GFP-Lmo7 mRNA ( approximately 25-50 %). (B-B") Representative image of the boundary between cell clusters with (asterisks) and without GFP-Lmo7. Cell outlines are marked by phalloidin staining. (C) Quantification of apical domain size. Control uninjected cells (n=96), *lmo7*-ATGMO+GFP cells (n=94) and *lmo7*-ATGMO+GFP-Lmo7 cells from more than five different embryos. Statistical significance of the difference between the median values was assessed by the Student's t-test.

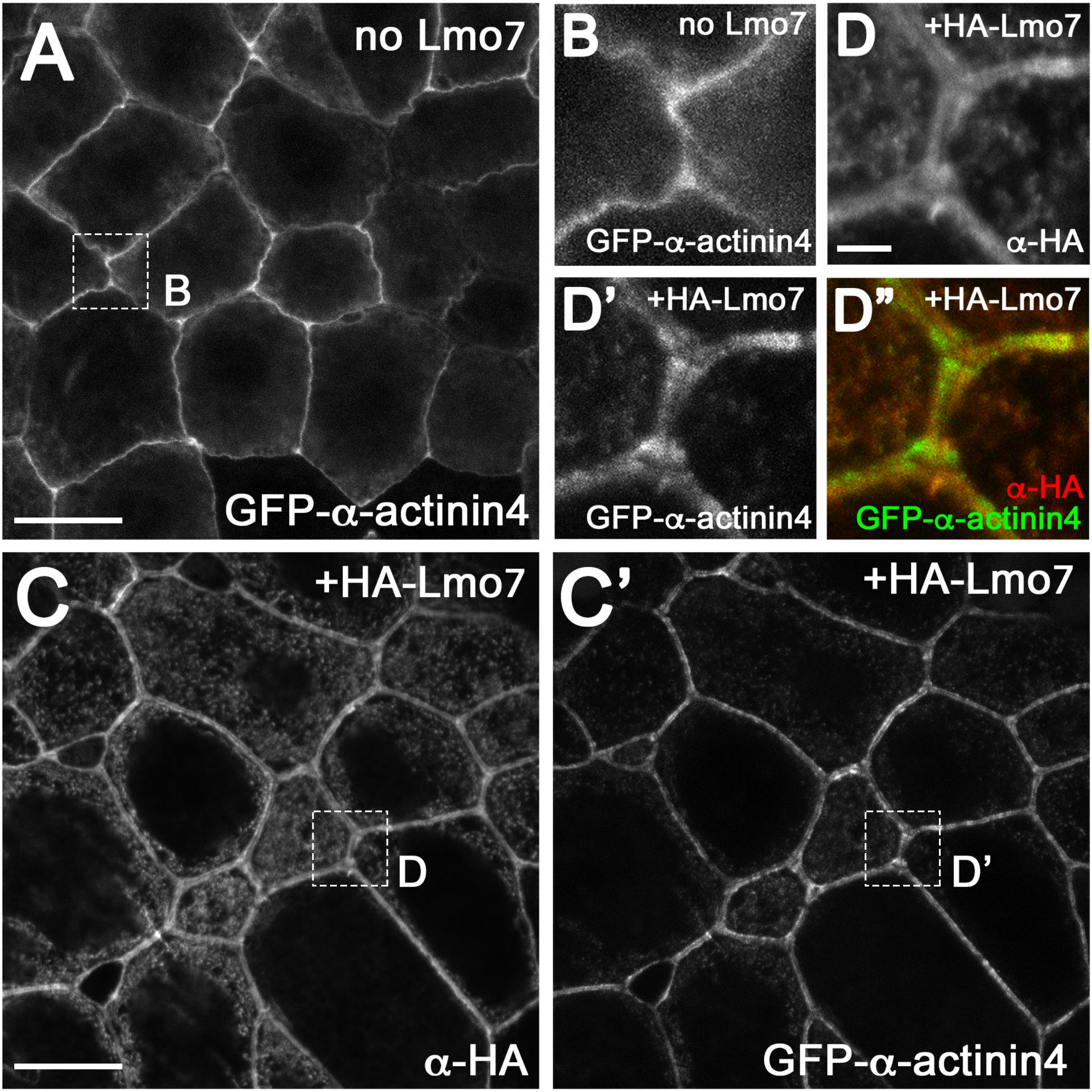
A
no Lmo7
B
GFP-α-actinin4
B
no Lmo7
GFP-α-actinin4
D
+HA-Lmo7
α-HA
D'
+HA-Lmo7
GFP-α-actinin4
D"
+HA-Lmo7
α-HA
GFP-α-actinin4
C
+HA-Lmo7
D
α-HA
C'
+HA-Lmo7
D'
GFP-α-actinin4

### Figure S2

**Supplemental Figure 2. Lmo7 promotes α-actinin enrichment in perijunctional actomyosin bundles**

mRNA encoding GFP-α-actinin-4 (200 pg) was injected into 4-8 cell stage embryos with or without mRNA encoding HA-Lmo8 (500 pg). (A, B) GFP-α-actinin-4 localizes at apical junctions and forms a single band. An area marked by a rectangle in A is enlarged in B. (C-D") HA-Lmo7 promotes GFP-α-actinin-4 association with apical junctions. Areas marked by rectangles in C-C' are enlarged in D-D". GFP-α-actinin-4 forms thick double bands that largely overlap HA-Lmo7. Scale bars: 10 μm in A and C. 2 μm in D.

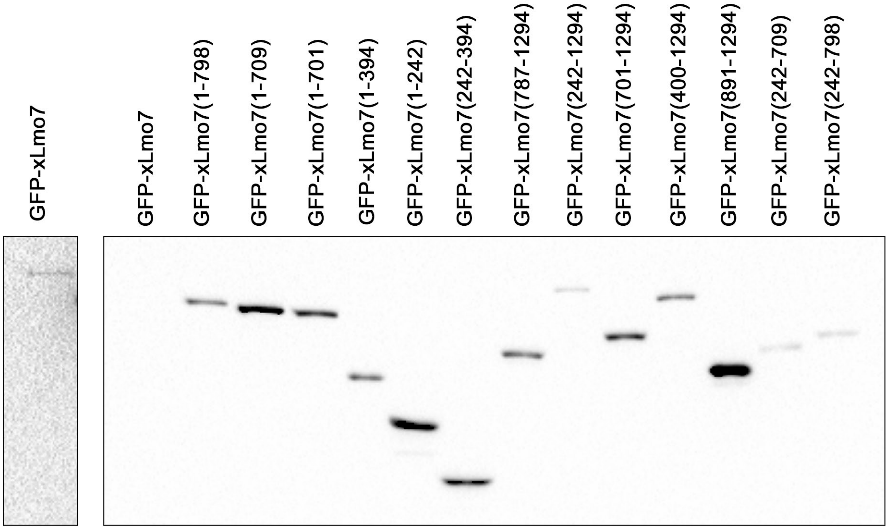
GFP-xLmo7
GFP-xLmo7
GFP-xLmo7(1-798)
GFP-xLmo7(1-709)
GFP-xLmo7(1-701)
GFP-xLmo7(1-394)
GFP-xLmo7(1-242)
GFP-xLmo7(242-394)
GFP-xLmo7(787-1294)
GFP-xLmo7(242-1294)
GFP-xLmo7(701-1294)
GFP-xLmo7(400-1294)
GFP-xLmo7(891-1294)
GFP-xLmo7(242-709)
GFP-xLmo7(242-798)

Figure S3

**Supplemental Figure 3. Expression levels of GFP-Lmo7 and its deletion mutants in *Xenopus* embryos**

GFP-Lmo7 construct mRNAs (1 ng) were injected into 4-8 cell stage embryos. Total embryo lysates were collected at stage 11. Expression levels of GFP-tagged Lmo7 constructs were assessed by immunoblotting with anti-GFP antibodies.

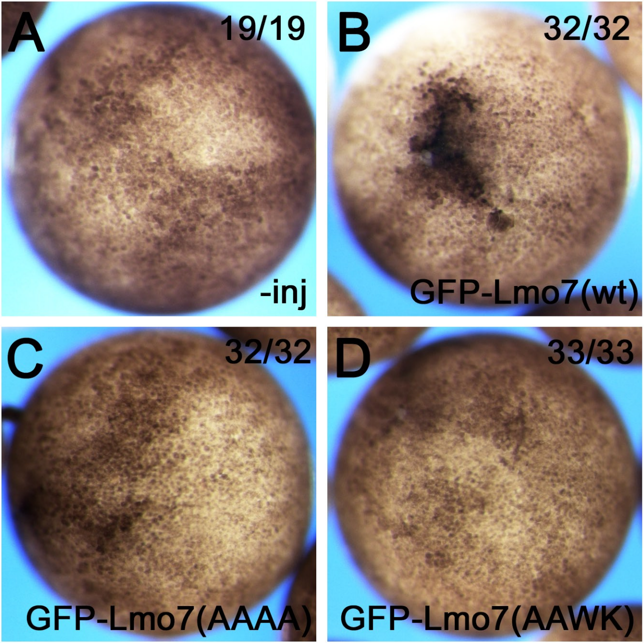
A
19/19
-inj
B
32/32
GFP-Lmo7(wt)
C
32/32
GFP-Lmo7(AAAA)
D
33/33
GFP-Lmo7(AAWK)

Figure S4

**Supplemental Figure 4. The DUF4757 domain is required for Lmo7-mediated apical constriction**

Embryos were injected into two blastomeres of 4-8 cell *Xenopus* embryos. Apical pigment granule accumulation was analyzed at stage 11. (A-D) Representative images with total numbers of embryos expressing the shown phenotype. (A) Uninjected control, (B) GFP-Lmo7, (C) GFP-Lmo7AAAA and (D) GFP-Lmo7AAWK. GFP-Lmo7AAAA and GFP-Lmo7AAWK carry substitutions in the conserved WK-WQ sequence in DUF4757.

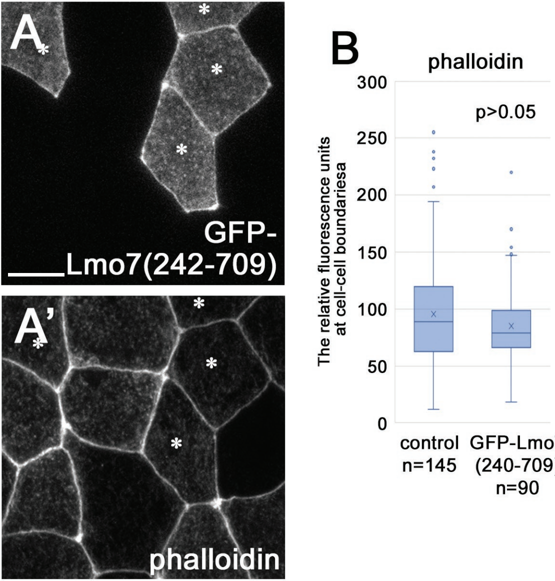
A
GFP-
Lmo7(242-709)
A'
phalloidin
B
phalloidin
300
250
200
150
100
50
0
p>0.05
The relative fluorescence units
at cell-cell boundariesa
control
n=145
GFP-Lmo7
(240-709)
n=90

### Figure S5

**Supplementary Figure 5. GFP-Lmo7(aa 242-709) does not induce F-actin accumulation at apical junctions**

GFP-Lmo7(aa 242-709) mRNA (1 ng) was injected into one blastomere of 4-8 cell stage embryos. Embryos are stained with phalloidin. (A, A') Representative images of embryos. Asterisks show cells expressing GFP-Lmo7(aa 242-709). (B) Quantification F-actin accumulation at apical junctions. Fluorescence intensity of phalloidin was measured at 3-10 locations within individual perijunctional F-actin bundles. Scale bar: 10 μm

Primer list

| | |
|---|---|
| xLmo7-2F-EcoRI | gaattcggaatggaatgaaaattc |
| xLmo7-242F-EcoRI | gaattccatgtcccatcgtagg |
| xLmo7-400F-EcoRI | gaattcggagagggaaacccg |
| xLmo7-700F-EcoRI | gaattcgtacagtgacctgagaat |
| xLmo7-787F-EcoRI | gaattctggcaagaatgactgg |
| xLmo7-898F-EcoRI | gaattcttgggatccagaaga |
| xLmo7-242R-stop-NheI | gctagctcagtcatctttgctgctg |
| xLmo7-394R-stop-NheI | gctagctcaggtttccctctcctgt |
| xLmo7-709R-stop-NheI | gctagctcaaggtttctggttaatgc |
| xLmo7-790R-stop-NheI | gctagctcagtcattcttgccatatc |
| xLmo7-1028R-stop-NheI | gctagctcattttaaaacattattcctttgtc |
| xLmo7-1274R-stop-NheI | gctagctcacatggaggttgg |
| xLmo7-WQWK(AAAA)-F | gagcaggatcagcagGCgGCgaatgatttagcaaaaGCgGCgaatcgtcgaaaaagc |
| xLmo7-WQWK(AAAA)-R | gctttttcgacgattcGCcGCttttgctaaatcattcGCcGCctgctgatcctgctc |

morpholino

| | |
|---|---|
| xLmo7 ATGMO | GAATTTTCATTCCATTCCATTG |

### Supplemental table 1
